# The *lac* operon in uropathogenic *Escherichia coli* enhances intracellular growth by enabling host glycan utilization

**DOI:** 10.64898/2025.12.03.691977

**Authors:** Jesus Bazan Villicana, Devina Puri, Taylor M. Nye, Rebecca L. Hurto, Jennie Hazen, Nathaniel C. Gualberto, Edward D. B. Lopatto, Jerome S. Pinkner, Karen W. Dodson, Lydia Freddolino, Scott J. Hultgren

## Abstract

The *lac* operon in *Escherichia coli* has been used as a model of gene regulation throughout biology since its characterization in the early 1960s. Despite the myriad of biotechnology applications that arose from characterization of the *lac* system, the explanation for the importance of a functional *lac* operon in normal bladder colonization by uropathogenic *E. coli* remains unknown given that the prototypical substrate, lactose, is not normally known to be readily available within the urinary tract. Here, we identified a unique uropathogenic clinical isolate (5.3r) that has a two codon deletion in the LacY permease, leading to impaired β-galactoside metabolism and attenuation of development of critical intracellular bacterial communities (IBCs) in which UPEC replicates to high numbers, and then disseminates during urinary tract infection. Further, we show that expression of a functional *lacY* permease gene is sufficient to rescue defects in IBC size and number in 5.3r. In addition, we demonstrate that UPEC are able to utilize the disaccharide galactose β-1,4 N-acetylglucosamine (LacNAc) – which appears as the terminal glycan in a subset of the glycoproteins that decorate the apical surface of bladder epithelial cells – as a sole carbon source in a *lac* system dependent manner. These data suggest that the *lac* operon is important to the growth and development of intracellular *E. coli* through metabolism of host bladder cell glycans.

## INTRODUCTION

Urinary tract infections (UTIs) are among the most common infectious diseases in the world, with nearly two-thirds of all women and 5% of men worldwide estimated to experience at least one UTI in their lifetime (1–6). In addition, UTIs can be highly recurrent (rUTI), where women with a past UTI have a 25% chance to develop rUTI within 6 months, in part due to epigenetic imprinting in the bladder mucosa upon the initial infections that predispose to recurrence (7, 8). Globally, rUTIs contribute to tremendous morbidity for millions of women, as well as thousands of preventable deaths, all of which underscore their enormous medical and economic burden (9–11). Uropathogenic *Escherichia coli* (UPEC) is the most common uropathogen, accounting for 75- 90% of community-acquired UTIs and over 50% of nosocomial UTIs worldwide (12, 13).

UPEC are extremely genetically and phenotypically diverse, with isolates differing by up to 40% in their accessory genes (14). The pangenome of *E. coli* contains upwards of 16,000 genes, of which individual strains carry approximately 3,000 core genes and 2,000 variable genes (15). Despite this genetic diversity, during uropathogenesis, most UPEC strains bind to and invade superficial bladder facet cells, where they form biofilm-like masses of >10^4^ cells referred to as intracellular bacteria communities (IBCs) (16–20). IBC formation progresses through specific developmental stages that support proliferation, persistence, and eventual dissemination to neighboring cells (16). These developmental stages include **i)** early: in the initial phase, single UPEC invade individual superficial umbrella cells of the bladder and rapidly multiply, forming a cluster of rod-shaped bacteria within the cytoplasm; **ii)** middle: the IBCs mature into densely arranged colonies of >10^4^ bacteria with biofilm characteristics; and **iii)** late: some bacteria at the outer edge of the IBC develop a filamentous morphology, facilitating their spread to neighboring cells where they then repeat the IBC cycle (16).

Several animal studies suggest that the IBC cycle allows UPEC to gain a foothold within the bladder, and bacteria that form the initial IBCs are the ones that survive to initiate the ongoing infection (21). Importantly, IBCs have also been detected in human urine of patients experiencing UPEC UTIs (17, 18). Thus, the IBC cycle is clinically relevant, and determining factors governing its formation and growth is of critical importance to understanding UPEC pathogenesis.

Through characterization in murine models of cystitis, previous studies have demonstrated that several genes associated with metabolism, biofilm formation, and stress response are upregulated in these bacteria proliferating intracellularly (**Table 1**). For instance, genes associated with iron acquisition play key roles in IBC development (22, 23). Additionally, there is strong evidence of UPEC switching to alternative sugar metabolism (e.g. galactose or sorbitol) while inside the host (22). A 2.5-fold increase in *lacZ* gene expression in IBCs was identified relative to a static laboratory culture, which strongly suggests a low glucose environment is encountered by UPEC in the bladder epithelium and that galactoside metabolism genes play an essential role in this intracellular pathogenesis cycle (22). Additionally, UPEC mutants lacking either *lacZ* or *galK* genes resulted in smaller IBCs, decreased ability to re-invade neighboring umbrella cells, attenuated colonization of the urinary tract, and were defective for maintenance of chronic cystitis (22).

**Table 1.**
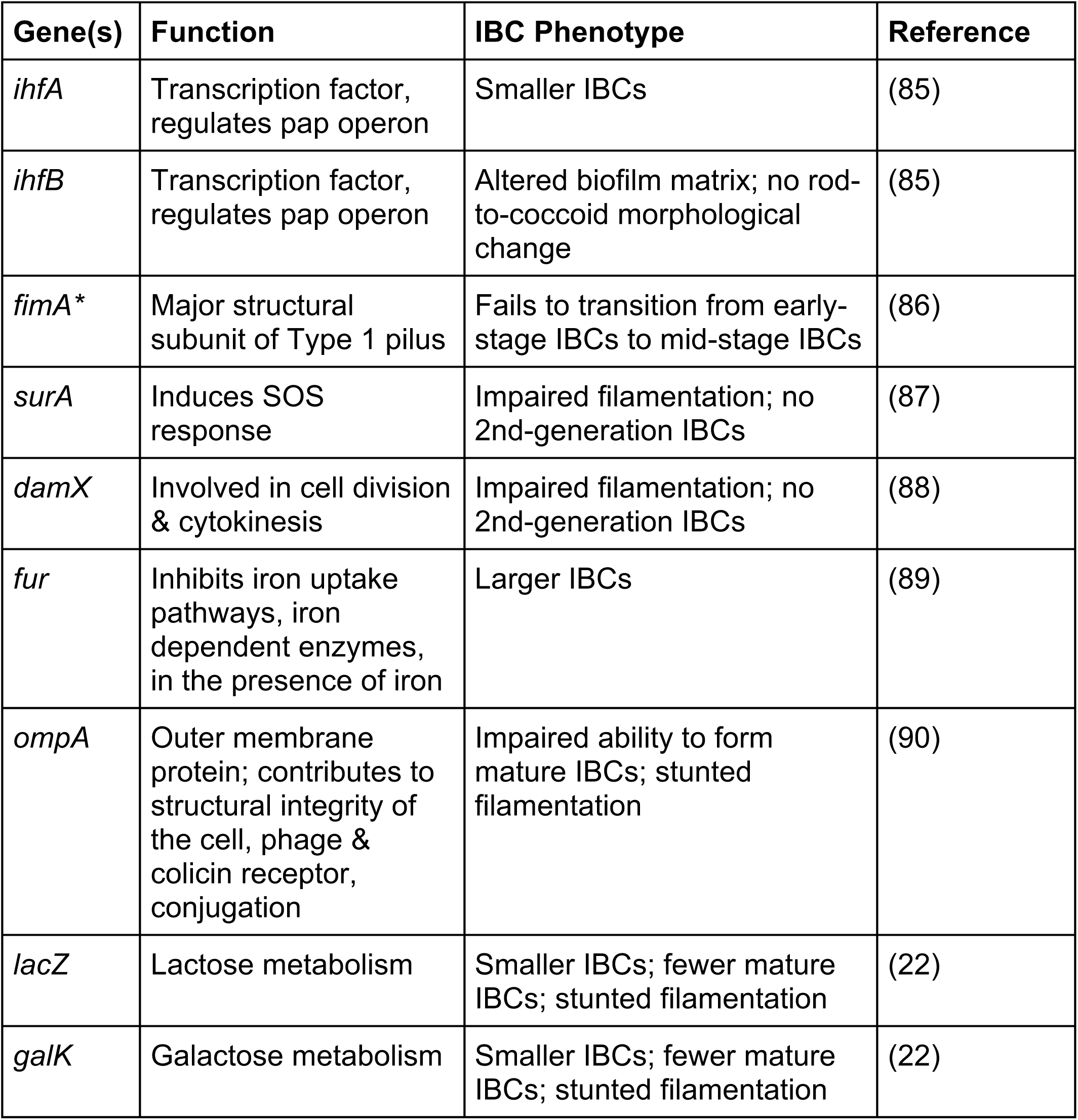
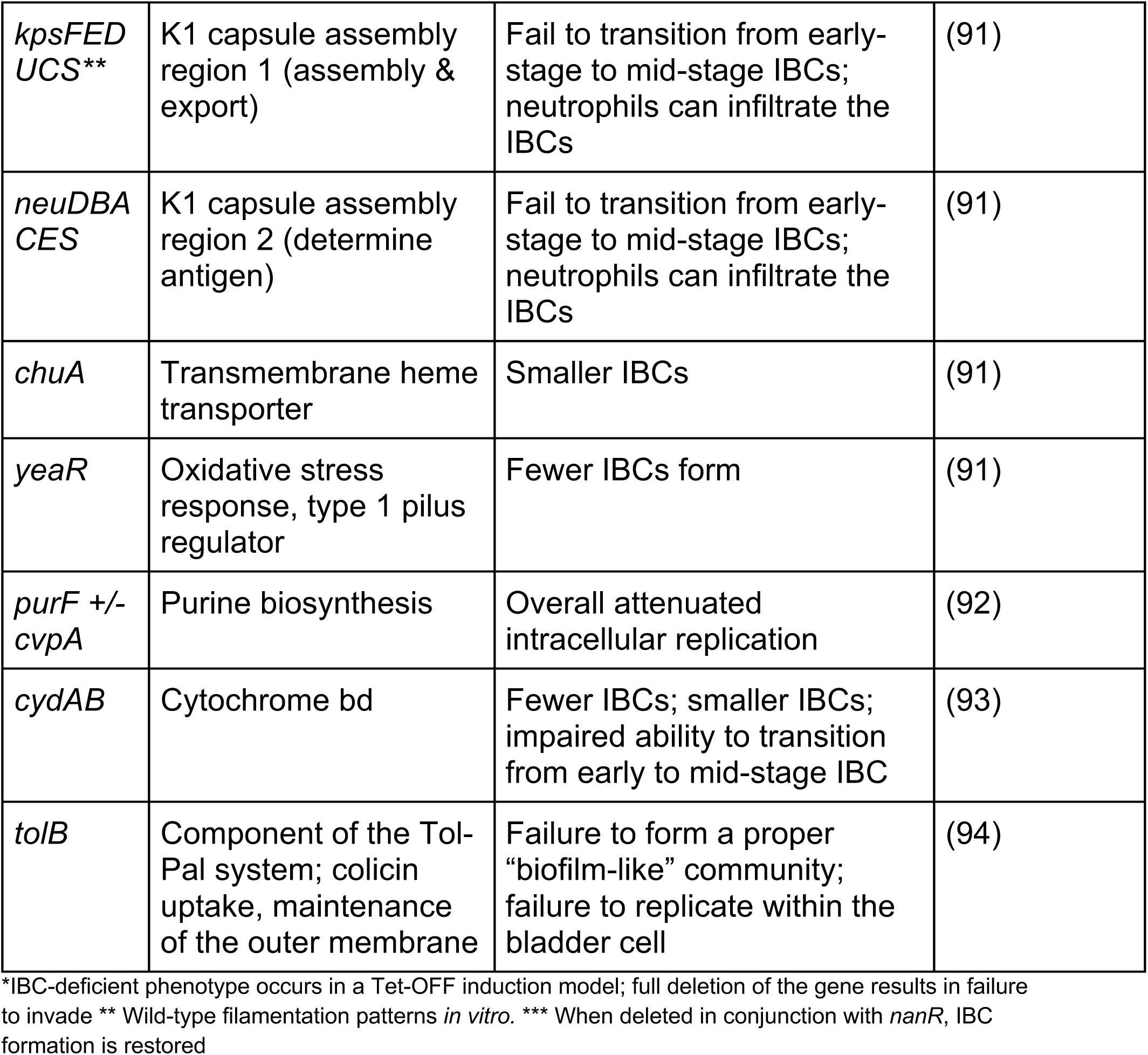
Genes with known roles in the formation of intracellular bacterial communities in UPEC. List of genes that when altered, are known to affect IBC morphology, IBC cycle kinetics, and/or IBC number. Not depicted are genes that result in complete lack of IBC formation.

| **Gene(s)** | **Function** | **IBC Phenotype** | **Reference** |
| --- | --- | --- | --- |
| *ihfA* | Transcription factor, regulates pap operon | Smaller IBCs | (85) |
| *ihfB* | Transcription factor, regulates pap operon | Altered biofilm matrix; no rod-to-coccoid morphological change | (85) |
| *fimA* * | Major structural subunit of Type 1 pilus | Fails to transition from early-stage IBCs to mid-stage IBCs | (86) |
| *surA* | Induces SOS response | Impaired filamentation; no 2nd-generation IBCs | (87) |
| *damX* | Involved in cell division & cytokinesis | Impaired filamentation; no 2nd-generation IBCs | (88) |
| *fur* | Inhibits iron uptake pathways, iron dependent enzymes, in the presence of iron | Larger IBCs | (89) |
| *ompA* | Outer membrane protein; contributes to structural integrity of the cell, phage & colicin receptor, conjugation | Impaired ability to form mature IBCs; stunted filamentation | (90) |
| *lacZ* | Lactose metabolism | Smaller IBCs; fewer mature IBCs; stunted filamentation | (22) |
| *galK* | Galactose metabolism | Smaller IBCs; fewer mature IBCs; stunted filamentation | (22) |
| *kpsFED*<br>*UCS* ** | K1 capsule assembly region 1 (assembly & export) | Fail to transition from early-stage to mid-stage IBCs; neutrophils can infiltrate the IBCs | (91) |
| *neuDBA*<br>*CES* | K1 capsule assembly region 2 (determine antigen) | Fail to transition from early-stage to mid-stage IBCs; neutrophils can infiltrate the IBCs | (91) |
| *chuA* | Transmembrane heme transporter | Smaller IBCs | (91) |
| *yeaR* | Oxidative stress response, type 1 pilus regulator | Fewer IBCs form | (91) |
| *purF +/-*<br>*cvpA* | Purine biosynthesis | Overall attenuated intracellular replication | (92) |
| *cydAB* | Cytochrome bd | Fewer IBCs; smaller IBCs; impaired ability to transition from early to mid-stage IBC | (93) |
| *tolB* | Component of the Tol-Pal system; colicin uptake, maintenance of the outer membrane | Failure to form a proper “biofilm-like” community; failure to replicate within the bladder cell | (94) |

\*IBC-deficient phenotype occurs in a Tet-OFF induction model; full deletion of the gene results in failure to invade \*\* Wild-type filamentation patterns *in vitro*. \*\*\* When deleted in conjunction with *nanR*, IBC formation is restored

To better understand the genetic pathways important to the development of robust IBCs across diverse UPEC isolates, we assessed IBC development in representative clinical strains across a 24 hour time-course in a murine cystitis model. Here, we highlight a unique clinical isolate, 5.3r, which has a two codon deletion in the *lac* operon lactose permease, *lacY*, that renders it unable to grow with lactose as the sole carbon source in culture and exhibits defects in IBC formation. We are able to restore lactose utilization and achieve prototypical IBCs by complementation with *lacY* from the model UPEC strain, UTI89. Additionally, we investigated the differences in the utilization of β-galactosides and similar carbon sources by 5.3r and UTI89, and found that the presence of the functional UTI89 *lacY* allele determines these growth differences. Finally, we show that UPEC (both UTI89 and, with appropriate *lacY* complementation, 5.3r) is able to metabolize the disaccharide galactose β-1,4 N-acetylglucosamine (LacNAc), which is a terminal disaccharide on a subset of the uroplakin glycans that coat the surface of bladder epithelial cells, in a *lac* operon dependent manner, providing evidence that UPEC may utilize novel host-derived carbon sources to establish UTIs within the bladder.

## RESULTS

### UPEC strain 5.3r is a lactose nonfermenting uropathogenic *E. coli* strain deficient in IBC formation

To better understand the genetic pathways that govern successful IBC development, we began by qualitatively screening a representative panel of eight genomically diverse strains, originally isolated from a prospective cohort of women with active UTIs, for their ability to form IBCs in our *in vivo* mouse model, maximizing representation of different UPEC clades. (14, 24). At 6 and 24 hours post infection (hpi) we inspected IBC formation by confocal microscopy. We found IBCs present in infected bladders with each strain tested, with the exception of clade A strains 2.1, 11.2p, and 11.3r. However, we observed significant differences in IBC cycle kinetics and morphologies between strains as summarized in **Table S1**. Strains 41.4p, 56.1a, and 20.1a, representing clades B1 and B2, showed similar IBC development to the prototypical clade B2 strain UTI89. However, the B2 strains 41.1a and 5.3r formed qualitatively smaller IBCs at 6 hpi and no IBCs were observed at 24 hpi, compared to the second generation IBCs that were observed for all other strains, suggesting that IBC development was impaired in these strains. Further, in 5.3r, filamentation was induced earlier than in UTI89 (6hpi), suggestive of an atypical IBC life-cycle compared to known prototypical strain behavior. We thus selected strain 5.3r for further study.

We previously identified 51 differentially expressed genes (DEGs) when comparing UTI89 in the inoculum to UTI89 recovered from the bladder at 6 hpi (22). Of these DEGs, 21 genes were positively regulated and 30 were negatively regulated. Thus, to understand 5.3r’s IBC phenotype, we began by analyzing the 5.3r sequences of these DEGs, and found nine DEGs with sequence differences between 5.3r and UTI89. Of these nine genes, four were in lambdoid prophage genes in multiple locations on the chromosomes, which are difficult to properly map and for which variations in sequence are likely simply due to different phage origins. Of the remaining five genes, only two, namely the lactose permease gene, *lacY,* and the poorly annotated DUF1971 domain-containing protein *yeaR*, had an indel between the strains; the other differences were point mutations **(Table S2)**. The 5.3r *lacY* (*lacY^5.3r^*) allele has a two codon deletion within the first transmembrane helix compared to the UTI89 *lacY* (*lacY^UTI89^*), suggesting that the *lacY^5.3r^* allele may be defective (**Fig 1A**) (25). The 5.3r *yeaR* allele contains a large N-terminal truncation which is also likely to substantially affect its function; however, given that the protein has not been heavily characterized (aside from having mutations arise in some specialized culture conditions, we focused first on the potential effects of the observed mutation in *lacY* (*26*). To test for potential impacts of the observed 5.3r *lacY* allele on biofilm formation, we assessed growth of 5.3r and UTI89 in M9 minimal media supplemented with lactose or either of the monosaccharides that comprise lactose (i.e., glucose and galactose) as the sole carbon source. While UTI89 showed robust growth in all three carbon sources, strain 5.3r could only grow on the monosaccharides glucose and galactose **(Fig 1B)**. Thus, we identified a clinical UPEC isolate, 5.3r, that is defective for the utilization of lactose as a sole carbon source.

**Figure 1:**
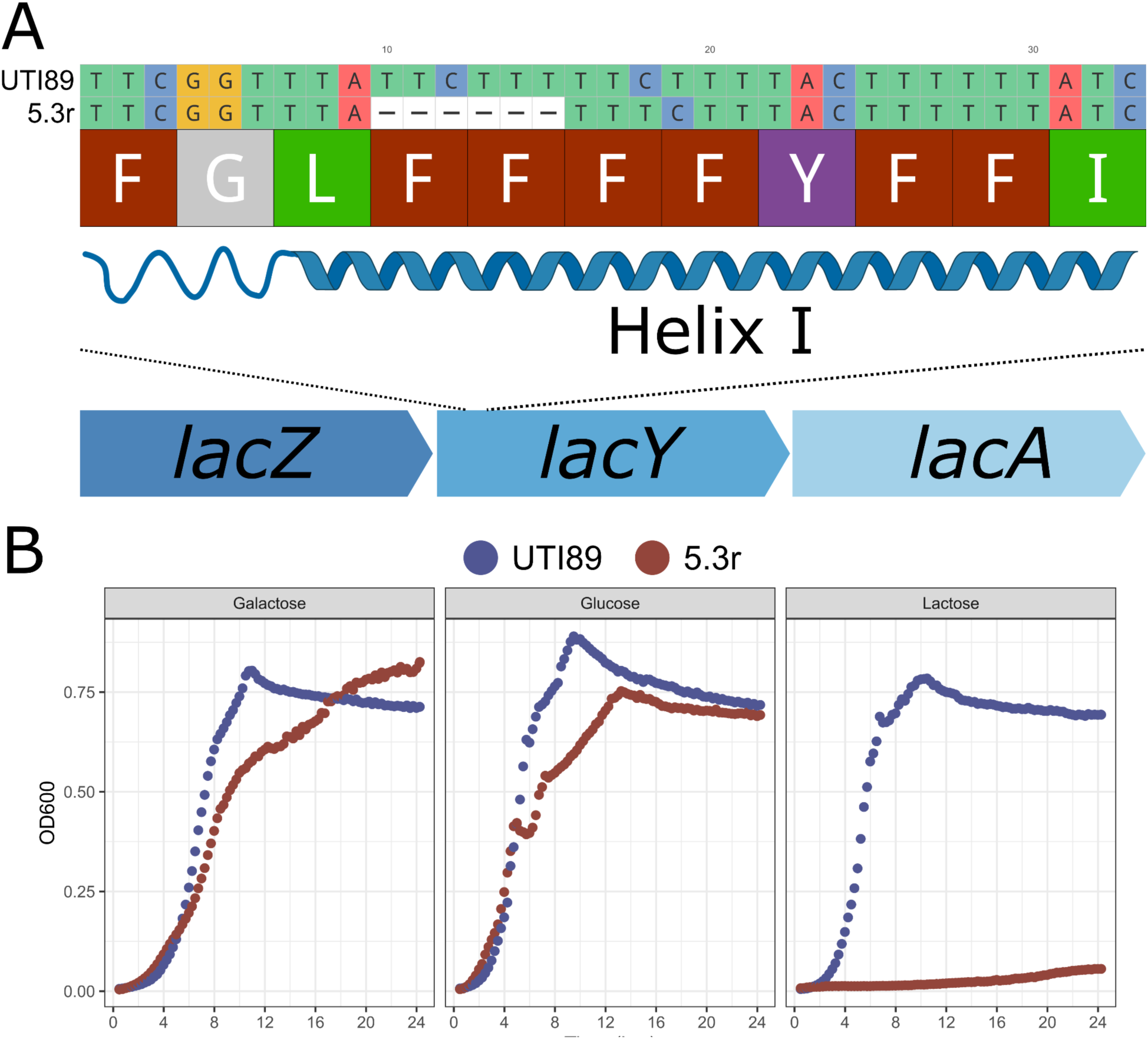
UPEC Strain 5.3r cannot use lactose as a sole carbon source *in vitro*. **(A)** Comparison of the *lacY* gene between UTI89 and 5.3r. (-) denotes absence of nucleotide at indicated position. **(B)** Growth of UTI89 and 5.3r in M9 minimal media with the indicated carbon sources including galactose, glucose, and lactose (0.4%). Plots represent the average OD600 from three biological replicates in UTI89 (navy) and 5.3r (copper).

### *lacY^UTI89^* but not *lacY^5.3r^* can complement 5.3r *in trans* to achieve growth in M9 lactose media

To determine if the identified two-codon deletion in the *lacY^5.3r^*was the reason for deficient growth of that strain in lactose-only media, we cloned the *lacY^UTI89^* into an IPTG inducible vector, pAb1, which also encodes a sfGFP gene (27). The resulting plasmid, pAB1-*lacY^UTI89^* (p*lacY^UTI89^*), was transformed into both 5.3r and UTI89 Δ*lacY* backgrounds. We also introduced a pAB1-*lacY^5.3r^* vector (p*lacY^5.3r^*), containing the two-codon deletion, into the 5.3r and UTI89 *ΔlacY* backgrounds to test if complementation with the *lacY^5.3r^* allele could restore the ability to utilize lactose as a sole carbon source in either strain. We also included a set of lactose non-fermenting strains, UTI89 *ΔlacY* and UTI89 *ΔlacZ*, for comparison. All strains were assayed for growth in M9 minimal media supplemented with glucose, glycerol, or lactose as sole carbon sources **(Fig 2).** All tested strains were able to grow with glucose and glycerol as sole carbon sources.

**Figure 2:**
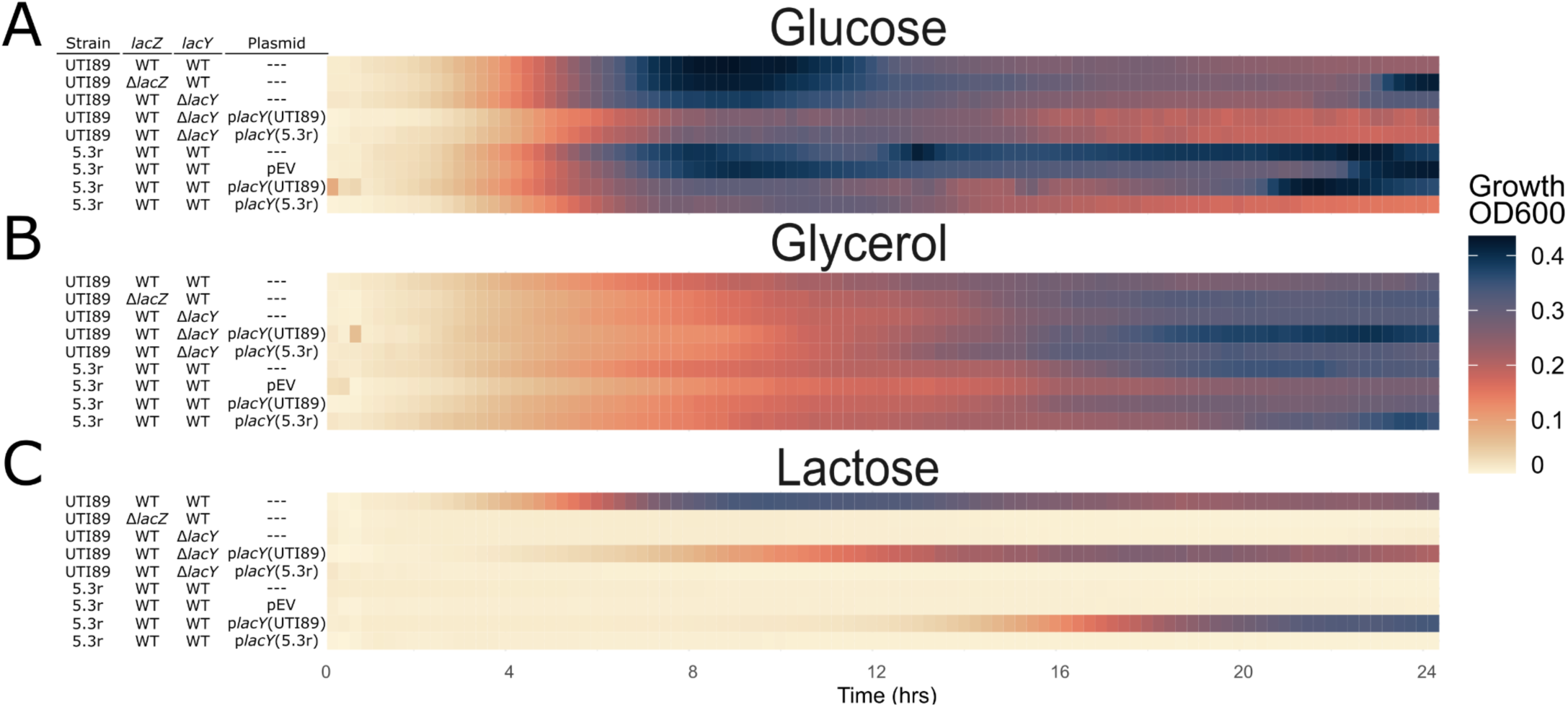
UTI89 *lacY* can complement 5.3r *in trans* to achieve growth in M9 lactose media. Heatmap of indicated strains grown in M9 media with 0.4% **(A)** glucose, **(B)** glycerol, or **(C)** lactose. Color gradient depicts increasing relative OD_600_ readings from white to black over time (hours).

However, while UTI89 could grow on lactose, strains 5.3r, UTI89 Δ*lacY*, and UTI89 Δ*lacZ* were all unable to grow in M9 minimal media containing lactose as a sole carbon source. Further, the deficiency in growth on lactose of 5.3r and UTI89 Δ*lacY* could be restored by complementation from p*lacY^UTI89^* but not p*lacY^5.3r^*demonstrating that the two codon deletion in 5.3r *lacY* results in a nonfunctional gene product that does not support growth in lactose as a sole carbon source.

### UPEC 5.3r exhibits attenuated IBC formation, which can be restored by a functional *lacY* gene

As strain 5.3r is both unable to utilize lactose as its sole carbon source, presumably due to an inability to import lactose, and exhibits atypical IBC morphology and developmental patterns (**Table S1**), we proceeded to further assess and quantify its impairment in IBC formation (16, 22). To do so, we compared IBC formation between UPEC strains 5.3r and UTI89 following transurethral inoculation and infection progression for 6 hours, a time at which the majority of infecting UTI89 are organized into IBCs within bladder epithelial cells (16, 22). Microscopic analysis of the bladders revealed that IBCs formed by strain 5.3r at this time point were defective relative to those in UTI89 (**Fig. 3A**), having decreased overall IBC area (65% reduced; **Fig. 3B**), IBC volume (77% reduced; **Fig. S1**), and IBC counts (82% reduced; **Fig. 3C**). Comparative analysis of 5.3r p*lacY^UTI89^* IBC structures at 6 hpi revealed IBCs that were similar in appearance to those generated by UTI89 and significantly larger than the ones formed by the original 5.3r strain (**Fig. 3A-B**). Additionally, 5.3r p*lacY^UTI89^* produced a higher number of IBCs compared to 5.3r (6.9-fold increase) and an overall increase in IBC area (2.6-fold increase; **Fig. 3B-C**). These findings collectively suggest that providing a functional *lacY* gene can alleviate the attenuation in IBC formation observed in UPEC strain 5.3r. Overall, these results suggest that the absence of lactose utilization in UPEC strain 5.3r leads to attenuation in IBC formation demonstrating at least one clinical strain case where a deficient *lacY* allele substantially alters the infection course of a UPEC strain.

**Figure 3.**
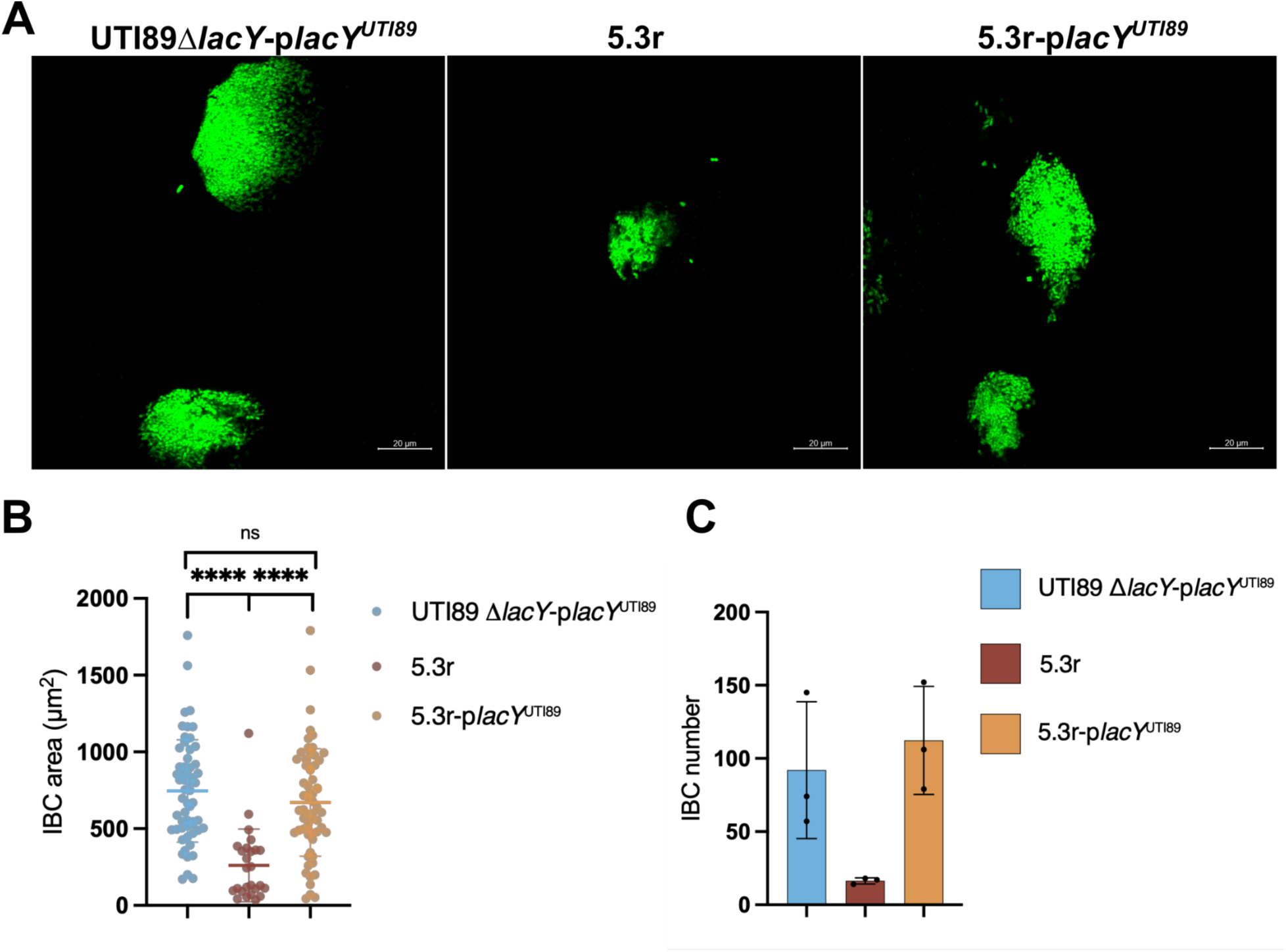
Functional *lacY* can restore IBC formation in UPEC 5.3r. **(A)** Representative micrographs of IBCs formed after 6 hours of infection with UTI89 Δ*lacY* p*lacY*^UTI89^ (left panel), 5.3r (middle panel), and 5.3r p*lacY*^UTI89^ (right panel). **(B)** Quantification of IBC area and **(C)** number after 6 hours of infection.

### Differential Carbon Utilization between UTI89 and 5.3r

Mammary epithelial cells are the only human cell types known to synthesize lactose, and lactose has only been detected in the urine of lactating mothers, making it possible that a carbon source other than lactose is being used to promote IBC development (28–30). Further, as the bladder contracts after voiding, epithelial cells shrink and large portions of the surface of the cell are internalized via endocytosis, allowing cell surface glycans that decorate uroplakins to enter the cells, possibly providing glycans for use by the growing IBC (31, 32). Thus, to better understand the function of LacY in the bladder epithelial cells, and how non-functional LacY^5.3r^ results in the development of fewer and smaller IBCs, we investigated differences in the ability of UTI89 and 5.3r to metabolize various carbon sources for growth. We utilized the Biolog microbial phenotype plates (PM1 and PM2A) with UTI89, 5.3r, and 5.3r p*lacY^UTI89^* to screen 192 carbon sources for lacY-dependent growth differences (**Fig. S2)**. In addition to lactose, only two of the screened carbon sources displayed a growth phenotype that was dependent on a functional *lacY* allele (**Fig. 4**): D-melibiose, a disaccharide of galactose and glucose with an ɑ-1,6 glycosidic linkage; and the monosaccharide D-methyl-α-galactoside, neither of which are known to be abundant in the bladder (33, 34). To expand our studies we considered glycans on the uroplakins, which coat the bladder cells, which might be utilized via the lac operon. A subset of mammalian uroplakins are terminated by the disaccharide galactose β-1,4 N-acetylglucosamine (LacNAc), including murine UP1b and bovine UPIII, which is the same glycosidic linkage hydrolyzed in lactose (galactose β-1,4 glucose) by the lac system (35, 36). As LacNAc is likely to be accessible to the *E. coli* cells when they form IBCs, and may thus become a substrate for LacZ, we tested whether UTI89 and 5.3r were able to utilize LacNAc as a sole carbon source. We found that UTI89 (but not 5.3r) can metabolize LacNAc in a *lac* operon dependent manner, requiring both *lacZ* and *lacY* for growth (**Fig. 5** **and Fig. S3**). Moreover, unlike lactose the presence of LacNAc alone is insufficient to trigger meaningful induction of the *lac* operon, requiring the addition of IPTG as an inducer under laboratory conditions in order to enable robust growth. Complementation of 5.3r with the *lacY^UTI89^* allele (but not an additional copy of the 5.3r allele) allows for growth in LacNAc.

**Figure 4:**
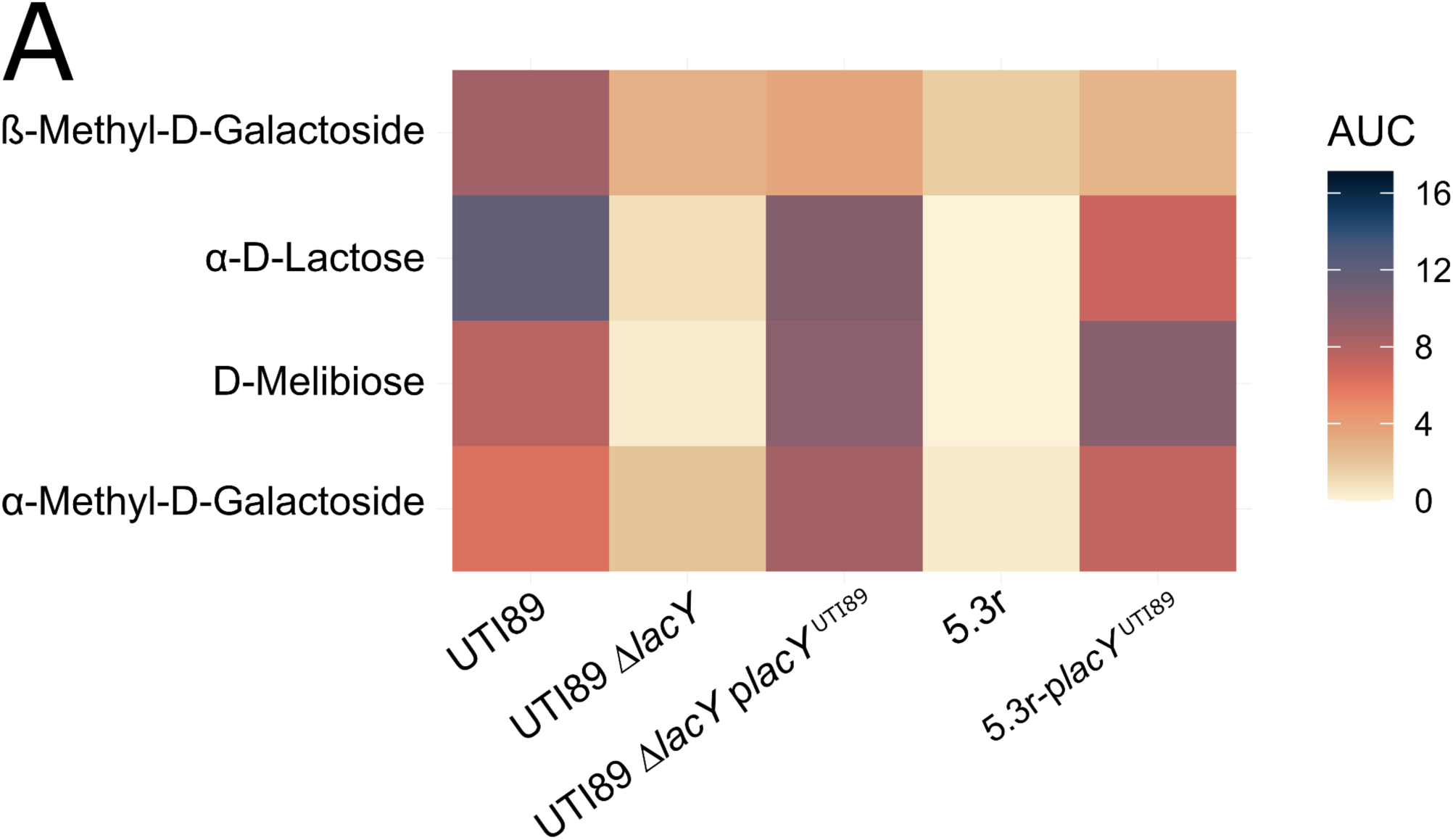
Differential utilization of carbon between UTI89 and 5.3r strains. **(A)** Heatmap depicting bacterial growth on selected compounds from the Biolog PM1 and PM2A carbon source plates between UTI89, UTI89 *ΔlacY,* UTI89 *ΔlacY* p*lacY*^UTI89^, 5.3r, and 5.3r p*lacY*^UTI89^. Color gradient depicts increasing relative AUC from OD600 readings from wheat to navy.

**Figure 5:**
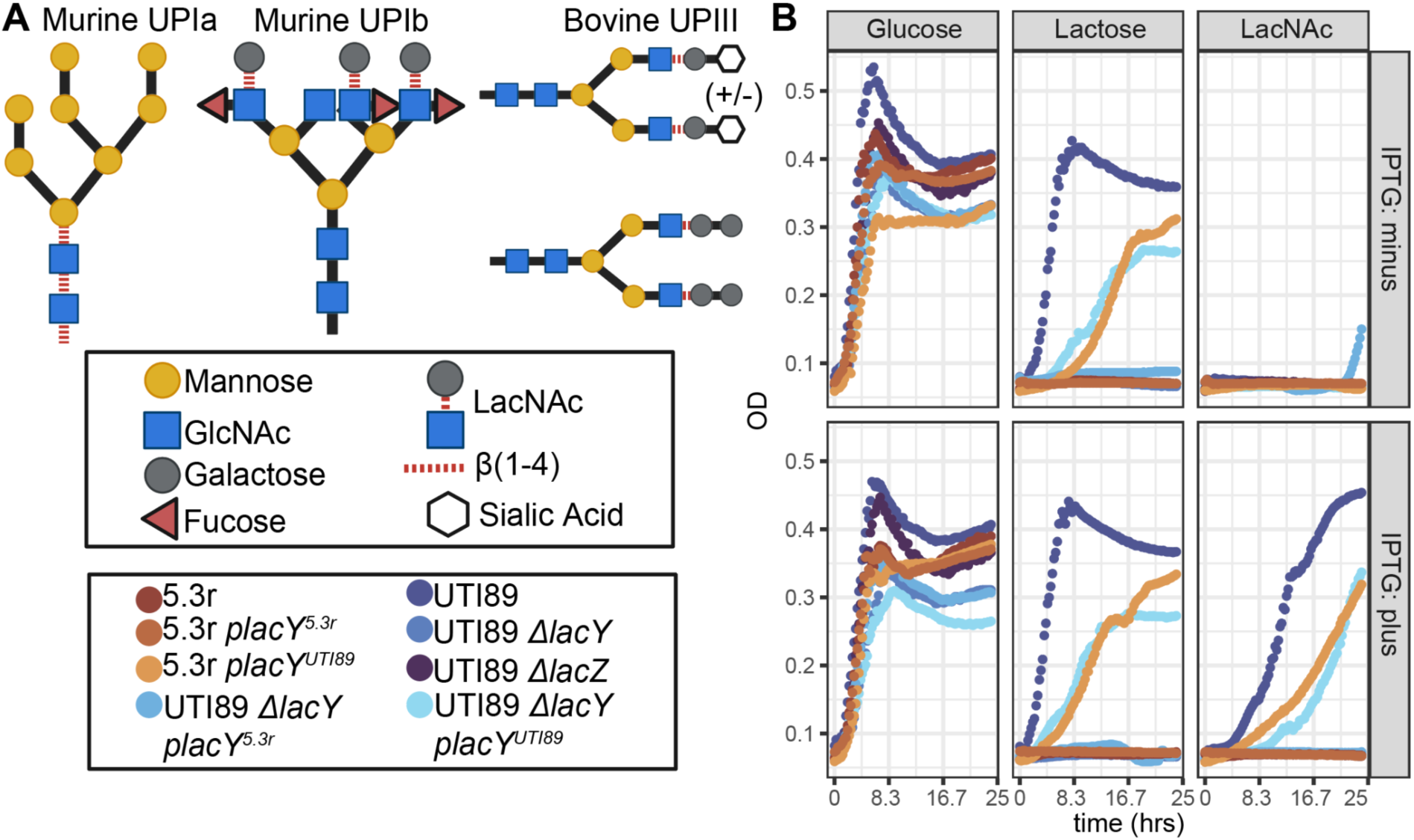
Mammalian bladder uroplakin structure and growth of strains in LacNAc. **(A)** Schema of dominant glycan structures that appear on murine uroplakin 1a and 1b and bovine uroplakin III (35, 36). LacNAc and known β-1,4 linkages that occur between other disaccharides are indicated by dashed red lines. **(B)** Representative growth curves depicting growth (OD_600_) of strains in M9 minimal media supplemented with either 0.4% glucose, lactose or LacNAc, in the presence and absence of the inducer IPTG.

## DISCUSSION

In this study, we examine the function of *lacY* within the context of clinical UPEC strains to better understand the role of the *lac* operon in a host environment. We have previously shown the importance of IBC formation for the onset and progression of UPEC UTIs and the upregulation of *lacZ* in IBCs (22). In addition, we have previously found that *Klebsiella pneumoniae,* which also can cause UTIs, also forms IBCs that show robust LacZ activity (44). In this work, we characterize a lactose non-fermenting UPEC clinical isolate, 5.3r, which exhibits an IBC-defective intracellular phenotype, similar to a UTI89 *ΔlacZ* mutant (22). We determined that the 5.3r *lacY* allele has a two codon deletion and thus 5.3r is incapable of using lactose as a sole carbon source. In addition, *lacY* from UTI89 (*lacY^UTI89^*) when expressed *in trans* in either UTI89 *ΔlacY* or 5.3r backgrounds showed *lacY*-dependent utilization of sugars including D-melibiose (a disaccharide of galactose and glucose with an α-1,6 glycosidic linkage) and the monosaccharide D-methyl-α-galactoside. Lastly, we show that N-acetyllactosamine (LacNAc) – a disaccharide composed of galactose and N-acetyglucosamine and a major component of mammalian glycans – is metabolized in a *lacY*-dependent fashion in the presence of the inducer IPTG. The atypical 5.3r *lacY* allele proved insufficient to support the LacNAc utilization in either a 5.3r or UTI89 background, whereas the UTI89 allele, which matches that found in the vast majority of sequenced *E. coli* strains, supported LacNac metabolism in either background. Taken together, these data suggest that *lacY* has a previously uncharacterized function in the transport of LacNAc that may help to establish UTIs in the bladder environment.

While 5.3r was a single clinical isolate with a nonfunctional *lacY* allele, lactose non-fermenting strains of *E.coli* have been well described and it is estimated that 5% of all clinical *E. coli* isolates are lactose non-fermenters (45, 46). This extends to urinary tract isolates, where upwards of 20% of isolates screened were lactose non-fermenters and exhibited increased rates of antimicrobial resistance (47–51). This suggests that lactose non-fermenting urinary tract isolates either have an alternative IBC cycle in human hosts or that additional host factors or comorbidities, e.g. being immunocompromised, permit these strains to cause UTIs.

The lac operon in *E. coli* has been extensively studied in laboratory conditions and encodes genes responsible for the metabolism of galactosides, including the most well-characterized galactoside, lactose (37–39). Previous studies have identified mutations within *lacY* that affect regulation of the *lac* operon or the transport or substrates, notably melibiose (40–43). Despite previous work with model UTI strains, the role of lactose metabolism genes in UPEC during the pathogenic IBC cycle is unclear. Our work raises important questions regarding the available carbon sources within the bladder during acute cystitis. Given that the lactose operon is canonically expressed when lactose is present and glucose is absent, it would be reasonable to assume that glucose levels must be low. In host cells, the previously observed induction of the *lac* system of *E. coli* may be accomplished by lactose/allolactose becoming available due to host cell degradation of uroplakins and/or other cellular components, or might arise by some parallel regulatory logic acting at the *lac* promoter that has not yet been characterized (22). The bladder cytosol is not known to contain high amounts of free lactose and previous work has implicated the importance of other carbon sources for UPEC to succeed within the bladder environment, including amino acids and short peptides (52–54). A possible explanation would be that non-lactose carbohydrates present in the bladder and within bladder urothelial cells can directly or indirectly stimulate expression of *lacZYA*. This would allow for utilization of degraded host factors such as LacNAc from glycosylated uroplakins, which are known to be important for UTI pathogenesis (55, 56).

It is known that complex sugar structures decorate superficial bladder cell surfaces, specifically uroplakin molecules (36, 55, 57). Many of these uroplakins are decorated with various sugars with known β1-4 glycosidic linkages on either terminal end, or present as structural scaffolding (58–60). UPEC are highly dependent on the type 1 pilus adhesin, FimH, binding to mannosylated uroplakins on the bladder surface to establish initial infection (61–65). Notably, molecular docking models predict that a mannose-(β1-4)-GlcNAc motif stabilizes fimH through interactions with the N-glycan core (62). Importantly, the presence of LacNAc has been found in human and murine urine (66–68). It is possible that a general moiety of mannose-(β1-4)-disacharride, whether it be LacNAc or another unknown glycan (**Fig. 5**), is important in the bladder for establishing an initial carbon source or inducing a unique metabolic state. Moreover, free glycans containing β1-4 glycosidic linkages have been detected in human urine, and could represent shedding of extracellular glycosylation from the bladder as it undergoes relatively high turnover during infections (68–70). Furthermore, when the bladder contracts after voiding, epithelial cells shrink and large portions of the cell are internalized via endocytosis, which could provide internalized glycans to be broken down by intracellular UPEC *in vivo* after initial attachment (31, 71–73).

Our results provide further evidence that galactoside metabolism genes impact UPEC’s intracellular pathogenic cascade in IBCs, given the two-codon difference in *lacY* between UTI89 and 5.3r and that expression of the *lacY^UTI89^* allele in 5.3r is sufficient to enable the UTI89 IBC and lactose metabolism phenotypes (22). Lactose metabolism is essential for UTI diagnosis as it is one of the methods for distinguishing *E. coli* from other Gram-negative bacteria in the urine, thus the prevalence of atypical, lactose non-fermenting strains may pose an additional diagnostic challenge (49). Recent work has also highlighted the diversity in regulation of the *lac* operon across *E.coli* species, which may further aid in understanding the importance of variant *lacY* alleles in regulatory and developmental pathways. (74). Hence, the understanding of lactose utilization pathways of non-fermenting UPEC strains is essential and understanding these metabolic processes could identify new therapeutic targets.

## METHODS

### Bacterial Strains and Growth Conditions

Experiments were conducted with the prototypical UPEC strain UTI89 and the clinical strains, including 5.3r, along with otherwise-isogenic mutants detailed in **Table S3** (**24, 75**). Standard M9 minimal media was used (76), supplemented with 1x BME Vitamins (Sigma-Aldrich; B6891) and 1x Trace Element solution (Sigma-Aldrich; MBD0056).

Where appropriate, the following antibiotic concentrations were used: kanamycin 50 μg ml^-1^, ampicillin 100 μg ml^-1^ and chloramphenicol 20 μg ml^-1^.

### Strain Construction

Electrocompetent cells were generated as previously described (77). Deletion mutations were constructed using the previously described λred recombinase method, using either pKD4 or PKD13 as a template (78, 79). Antibiotic cassettes were removed using pCP20 and expressing the FLP recombinase (80). For all *lacY* ectopic expression constructs, the IPTG inducible plasmid, pAB1 (Addgene plasmid # 62547). Plasmid construction was performed using restriction enzyme cloning. All primers utilized in this study are listed in **Table S3**.

### Growth Curves

Generally, strains were grown overnight in shaking conditions at 37°C, in M9 media containing 0.2% glycerol for 16 hours. Strains were subsequently washed with PBS twice and diluted to an OD_600_ of 1 that was used to inoculate wells 1:100. Bacteria were cultured in a Corning 96-well flat bottom plate containing 150 µL of M9 minimal media with appropriate carbon sources at 0.4% w/v. Each well was covered with 100 µL of mineral oil for a total well volume of 250 µL, preventing evaporation. The plate was read using a Biotek H1 plate reader (settings: 37°C, shaking orbital, OD_600_ every 15 minutes), over a period of 24-48 hours. For glycerol-depleted growth curves, strains were grown in 0.02% glycerol overnight for 16 hours and 5 µL of culture was used to directly inoculate 145 µL of M9 media with appropriate carbon sources. For Biolog growth assays, cultures were washed twice with PBS and then diluted to a suspension of OD of 0.01 in M9 minimal media without a carbon source. Then 125 µL of the suspension was loaded in each well and read at OD_590_ every 15 minutes with orbital shaking over 24 hours. Data shown is the average of two Biolog plates per strain. For the growth curves in LacNAc, 3 mL overnight cultures were grown in 14 mL culture tubes at 37°C with shaking in M9 media (M9 salts, 2mM MgSO_4_,100 µM CaCl_2_, MOPS micronutrients, 1 µM ferric citrate) supplemented with BME Vitamins (Sigma-Aldrich; B6891) and 0.2% glucose. The overnight cultures were diluted (5 µL) into 145 µL of M9 media supplemented with BME vitamins and 0.4% carbon source (glucose, lactose, or LacNAc) with and without IPTG at 0.1 mM in a 96-well plate. To prevent evaporation, 100 µL of sterile mineral oil was gently added to each well. The OD_600_ values for each well were measured every 15 minutes over a 24-hour period while the cells were incubated at 37°C with orbital shaking in a BioTek Epoch 2 Microplate Spectrophotometer .

### Murine infection and sample preparation

Infections were performed in accordance with the guidelines for Institutional Animal Care and Use Committee (IACUC) at Washington University School of Medicine in St. Louis, MO. UPEC strains were cultured in LB at 37°C for 24 hours under static conditions, subsequently diluted 1:1,000 in fresh LB and grown for an additional 18-24 hours. 0.1 mM IPTG was added to UPEC containing the IPTG inducible plasmid pAB1, with or without the cloned *lacY^UTI89^*, after 3-4 hours of growth of the subculture. These cultures were washed and resuspended phosphate buffered saline (PBS), and 10^8^ cfu of UPEC were injected transurethrally into lightly anesthetized female 7–8 week-old C57BL/6 mice (Charles River Labs) in a 50 µl dose (81, 82). Anesthesia was provided via inhalation of 4% isoflurane. The UPEC infection was allowed to progress for 6 hours. Post-infection, mice were euthanized, and their bladders were aseptically removed.

Bladders were hemisected and placed on 6-well plates prefilled with PBS and containing Sylgard 184, Silicone Elastomer Kit (Dow Corning) on their base. With the lumen side facing up, bladder corners were pinned onto the elastomer and stretched. Splayed bladders were fixed in 4% paraformaldehyde, stained with 4’,6-diamidino-2-phenylindole (DAPI), and mounted on a glass slide using ProLong Gold Antifade Reagent (Invitrogen) for microscopy analysis. Screening of IBCs of the clinical isolates was performed by transurethral inoculation of 10^7^ cfu in a 50 µl dose, where each isolate was transformed with the pANT4 plasmid, a high copy number plasmid that constitutively expresses GFP under the tight regulation from a synthetic promoter (75, 83, 84).

### Microscopy and IBC analysis

Microscopy was performed at the Molecular Microbiology Imaging Facility at Washington University School of Medicine in St. Louis. Prepared bladder slides were imaged using Zeiss LSM880 Confocal Laser Scanning Microscope with Airyscan (Carl Zeiss Inc.) and Zeiss Axio Imager M2 (Carl Zeiss Inc.) for fluorescence microscopy. Images were acquired and exported using ZEN software (Carl Zeiss Inc.). To quantify IBC size, raw images from the green-fluorescent channel were imported into ImageJ software (NIH) for quantitative analysis. Intensity-based segmentation using in-built functions was employed to isolate the IBC region and create binary masks.

Quantification of size was performed using the “Analyze Particles” function, where the measurement parameters were set to include object area. All measurements were exported for statistical analysis using GraphPad Prism (GraphPad software). Volumetric analysis of IBCs was performed by processing confocal microscopy images using Volocity software (PerkinElmer).

## ACKNOWLEDGMENTS

We acknowledge the members of the Hultgren and Freddolino labs for their support of this research. We thank Dr. Matthew Chapman for his feedback on the manuscript. This work was funded by NIH grants R01 DK051406 (S.J.H.) and RO1 AI13467801 (S.J.H. and L.F.).

## Notes

### Competing Interest Statement

The authors have declared no competing interest.

