## Supplementary Tables for "The *lac* operon in uropathogenic *Escherichia coli* enhances intracellular growth by enabling host glycan utilization"

### S1 Table of IBC Phenotypes of selected strains

| Strain | Clade | Image | 6hpi* | Presence of 2nd gen IBCs at 24hpi* |
|---|---|---|---|---|
| UTI89 | B2 | 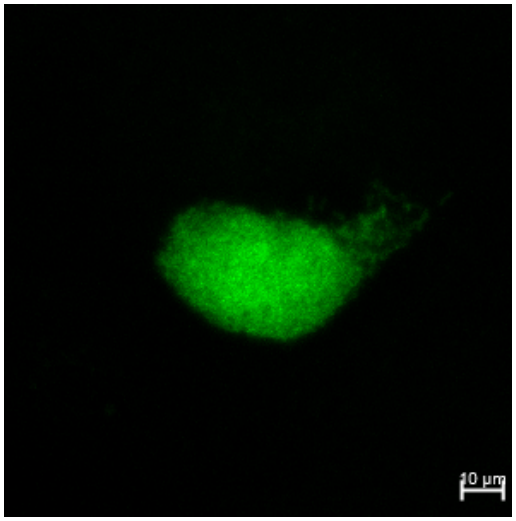  | Mid stage IBCs | Present[1] |
| 2.1a | A | N/A | No IBCs | No IBCs |
| 11.2p | A | N/A | No IBCs | No IBCs |
| 11.3r | A | N/A | No IBCs | No IBCs |
| 5.3r | B2 | 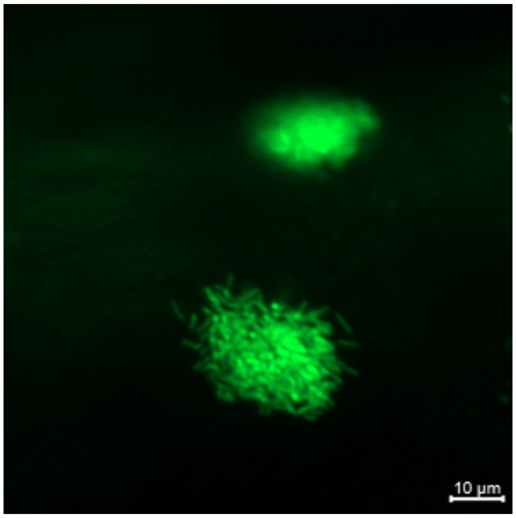  | Mid stage IBCs, filaments begin to form | No IBCs |

| | | | | |
|---|---|---|---|---|
| 20.1a | B2 | 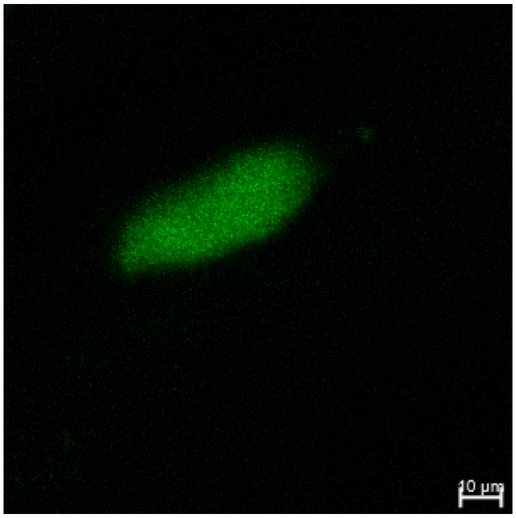 | Mid stage IBCs | Present |
| 41.1a | B2 | 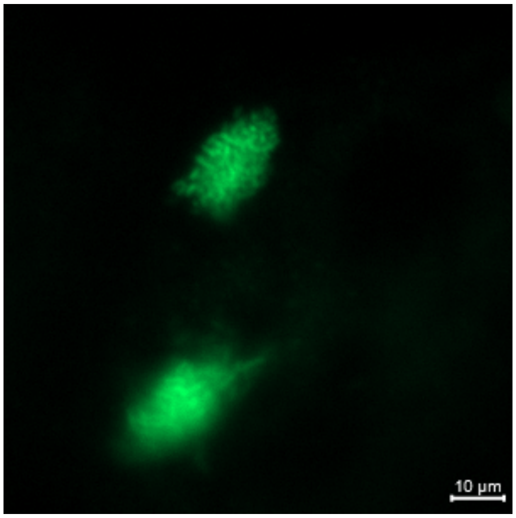 | Small mid stage IBCs | No IBCs |
| 41.4p | B1 | 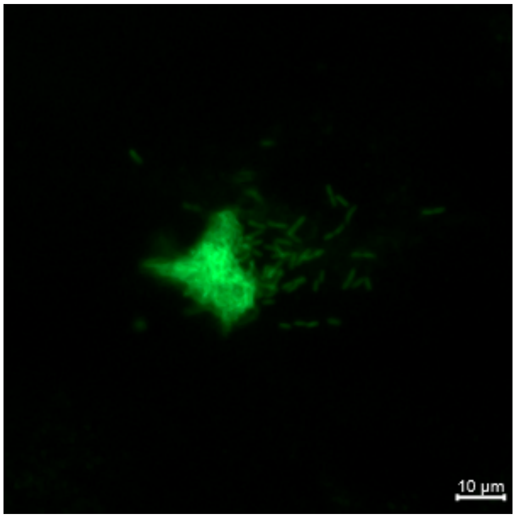 | Mid stage IBCs, resemble early stage UTI89 IBCs | No IBCs |

| | | | | |
|---|---|---|---|---|
| 56.1a | B1 | 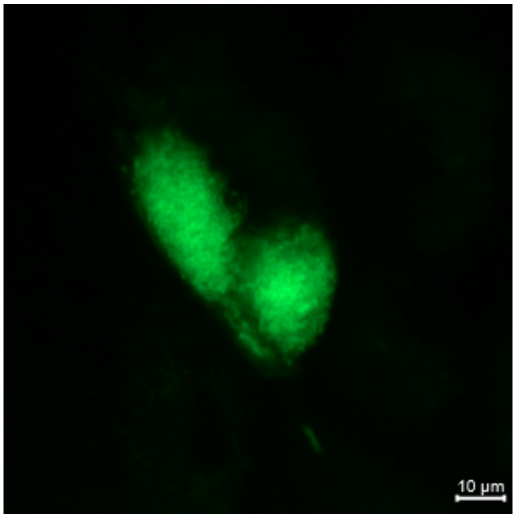 | Large mid stage IBCs | Present |

Shown above are individual timelines of IBC development for each strain with qualitative descriptions. Second generation IBCs as defined by (16, 95)

*hpi=Hours post infection

### S2 Table of genomic differences between UTI89 and 5.3r in IBC-related genes

| **locus** | **gene** | **% ID** | **Length (nt)** | **mutation** |
|---|---|---|---|---|
| UTI89_C0370(-) | *lacY* | 99.52 | 1248/1254 | delta_16-17 |
| UTI89_C1121(-) | *iroC* | 99.75 | 3660/3660 | T1034A |
| UTI89_C1338(-) | *sitB* | 99.64 | 828/828 | S32G |
| UTI89_C1995(-) | *yeaR* | 100 | 237/360 | N-terminal truncation |
| UTI89_C2180(-) | (Putative ABC transporter) | 99.83 | 1803/1803 | A202V |
| UTI89_C5111(+) | S-like lysis protein (prophage) | 95.83 | 216/216 | - |
| UTI89_C5112(+) | Lysozyme (prophage) | 63.23 | 462/462 | - |
| UTI89_C5113(+) | Endopeptidase (prophage) | 68.83 | 459/468 | - |
| UTI89_C5114(+) | Phage gene 65 (prophage) | 98.04 | 153/153 | - |

### S3 Table of strains and PCR primers used in this study

| ***Escherichia coli* (*E.coli*) strains** | | | |
|---|---|---|---|
| Strain Number | Genotype | Antibiotic Resistance | Reference or source |
| SJH-2 | UTI89 | None | (75) |
| SJH-18 | 5.3r | None | (24) |
| SJH-15 | 2.1a | None | |
| SJH-175 | 11.2p | None | |
| SJH-22 | 11.3r | None | |
| SJH-31 | 20.1a | None | |
| SJH-298 | 41.1a | None | |
| SJH-303 | 41.4p | None | |
| SJH-338 | 56.1a | None | |
| SJH-3253 | UTI89 Δ*lacZ* | $Amp^R$ | (22) |
| SJH-3941 | UTI89 HK::*kan* Δ*lacY* | $Kan^R$ | This Study |
| SJH-3943 | UTI89 HK::*kan* Δ*lacY* pAB1 | $Kan^R$,$Amp^R$ | This Study |
| SJH-3944 | UTI89 HK::*kan* Δ*lacY* pAB1-*lacY*$^{UTI89}$ | $Kan^R$,$Amp^R$ | This Study |

| | | | |
|---|---|---|---|
| SJH-3965 | UTI89 HK::*kan* Δ*lacY* pAB1-*lacY*$^{5.3r}$ | Kan$^R$,Amp$^R$ | This Study |
| SJH-3851 | 5.3r pAB1 | Amp$^R$ | This Study |
| SJH-3868 | 5.3r pAB1-*lacY*$^{UTI89}$ | Amp$^R$ | This Study |
| SJH-3966 | 5.3r pAB1-*lacY*$^{5.3r}$ | Amp$^R$ | This Study |

| **Primers used for strain construction** | | | |
|---|---|---|---|
| **Primer** | **Sequence (5' to 3')** | **Template** | **Amplicon Product** |
| JBV54 | cgatcgtctagaatgtactatttaaaaaac | UTI89 OR 5.3r | pAB1-lacY XbaI flanks |
| JBV55 | cgatcttctagattaagcgacttcattcac | | |
| JBV121 | atcaacttctgatggatgcga | UTI89 | kan-Δ*lacY* sequencing |
| JBV122 | cgctacagccaacaacaactg | UTI89 | kan-Δ*lacY* sequencing |
| JBV123 | ctgatatgttggtcggataaggcgctcgcgccgcatccgacattgattgcgtgtaggctggagctgctt | UTI89 | Δ*lacY* KO cassette |
| JBV124 | aataaccgggcaggccatgtctgcccgtatttcgcgtaaggaaatccattcatatgaatatcctccttag | | |

Antibiotic resistance markers: KanR, kanamycin; CmR, chloramphenicol; AmpR, ampicillin.
