## Supplementary Figures for "The *lac* operon in uropathogenic *Escherichia coli* enhances intracellular growth by enabling host glycan utilization"

### Supplemental Figure S1

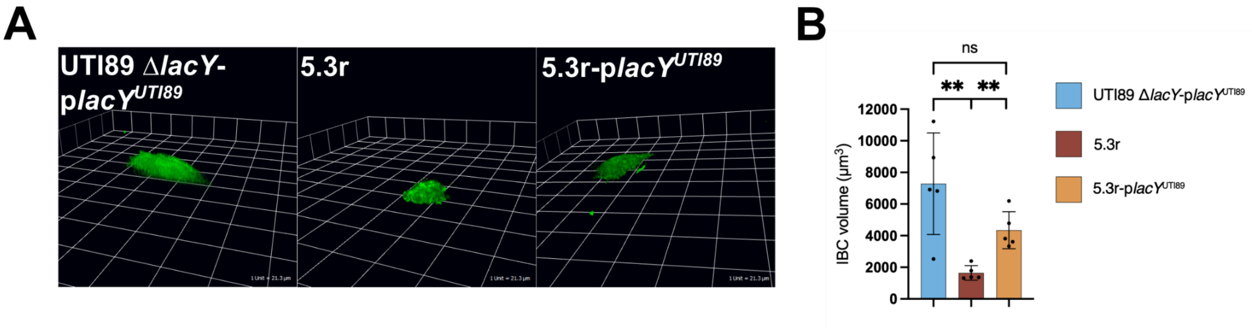

**(A)** 3D representation of IBCs formed after 6 hours of infection with (left panel) UTI89 Δ*lacY* p*lacY*$^{\mathrm{UTI89}}$, (middle panel) 5.3r, and (right panel) 5.3r- p*lacY*$^{UTI89}$. **(B)** Quantification of volume from rendered 3D images of IBCs formed after 6 hours of infection with UTI89 Δ*lacY* p*lacY*$^{\mathrm{UTI89}}$, 5.3r, and 5.3r p*lacY*$^{UTI89}$.

### Supplemental Figure S2

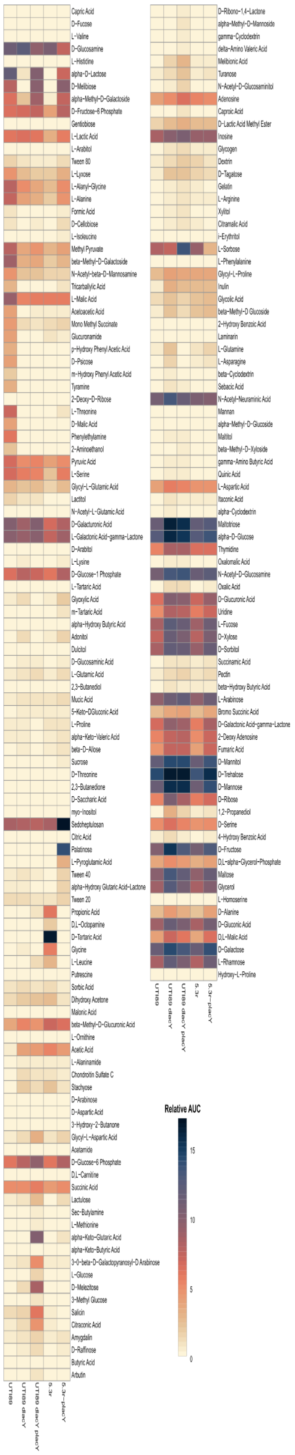

Heatmap depicting bacterial growth on Biolog PM1 and PM2A carbon source plates between UTI89, UTI89 *ΔlacY*, UTI89 *ΔlacY* p*lacY*$^{UTI89}$, 5.3r, and 5.3r p*lacY*$^{UTI89}$. Color gradient depicts increasing relative AUC from $OD_{600}$ readings from wheat to navy.

### Supplemental Figure S3

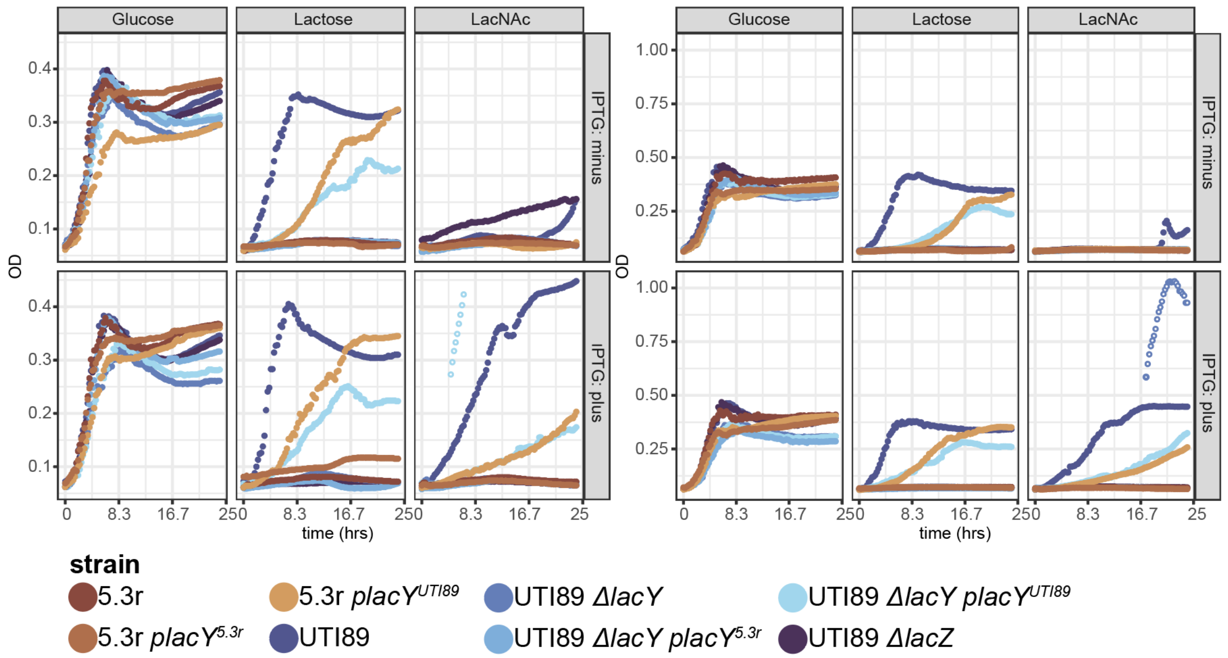

**Structure of bladder uroplakin structure and growth of strains in LacNAc.(A)** Schema of glycan structures that appear on indicated uroplakins. LacNAc and known β-1,4 linkages that occur between other disaccharides are indicated. **(B)** Growth curves depicting growth of strains in M9 media supplemented with either 0.4% glucose, lactose or LacNAc, with and without IPTG. Open circles indicate time points where $OD_{600}$ measurements were suspected to be impacted by bubble formation.
